# Association of polymorphisms of IL-6 pathway genes (*IL6, IL6R* and *IL6ST*) with COVID-19 severity in an Amazonian population

**DOI:** 10.1101/2022.09.21.508870

**Authors:** Fabíola Brasil Barbosa Rodrigues, Rosilene da Silva, Erika Ferreira dos Santos, Mioni Thieli Figueiredo Magalhães de Brito, Andréa Luciana Soares da Silva, Mauro de Meira Leite, Flavia Póvoa da Costa, Maria de Nazaré do Socorro de Almeida Viana, Kevin Matheus Lima de Sarges, Marcos Henrique Damasceno Cantanhede, Adriana de Oliveira Lameira Veríssimo, Mayara da Silva Carvalho, Daniele Freitas Henriques, Carla Pinheiro da Silva, Juliana Abreu Lima Nunes, Iran Barros Costa, Giselle Maria Rachid Viana, Maria Alice Freitas Queiroz, Sandra Souza Lima, Jeferson da Costa Lopes, Maria Karoliny da Silva Torres, Carlos David Araújo Bichara, Izaura Maria Vieira Cayres Vallinoto, Antonio Carlos Rosario Vallinoto, Eduardo José Melo dos Santos

## Abstract

Interleukin-6 have been recognized as a major role player in COVID-19 severity, being an important regulator of cytokine storm. Hence, the evaluation of the influence of polymorphisms in key genes of IL-6 pathway, namely *IL6, IL6R* and *IL6ST*, may provide valuable prognostic/predictive biomarkers on COVID-19. The present cross-sectional study genotyped three Single Nucleotide Polymorphisms - SNPs (rs1800795, rs2228145 and rs7730934) at *IL6, IL6R* and *IL6ST* genes, respectively, in 227 COVID-19 patients (132 hospitalized and 95 non-hospitalized). Genotype frequencies were compared between these groups. As control group, published data on gene and genotype frequencies was gathered from published studies from before the pandemic started. Our major results point to an association of *IL6* C allele with COVID-19 severity. Moreover, IL-6 plasmatic levels were higher among *IL6* CC genotype carriers. Additionally, the frequency of symptoms was higher at *IL6* CC and *IL6R* CC genotypes. In conclusion the data suggest an important role of *IL6* C allele and *IL6R* CC genotype on COVID-19 severity, in agreement with indirect evidences from literature about association of these genotypes with mortality rates, pneumonia, heightening of protein plasmatic levels proinflammatory driven effects.

## Introduction

The most striking characteristic of severe COVID-19 is the cytokine storm, where systemic inflammatory pathways are strongly activated. Along with TNF-alpha and IFN-gamma, high levels of IL-6 became one of the most important prognostic biomarkers in COVID-19 [1].

IL-6 is a multifunctional cytokine with both pro- and anti-inflammatory properties. IL-6 receptor bounded to membrane of hepatocytes and some leucocyte subpopulations is known as classic pathway, activating intracellular signaling pathways like JAK/STAT and being the gp130, encoded by *IL6ST* gene, responsible by the signal transduction. A second pathway, called trans-signaling, is constituted by the binding of IL-6 with a soluble receptor (sIL-6R), whose complex binds to soluble gp130 forms. The interplay between the pro-inflammatory properties of trans-signaling pathway and the anti-inflammatory signaling is highly regulated in all inflammatory processes [2] and seems to occupy a seminal position in COVID-19.

Additional evidence of the relevance of IL-6 and their receptors in COVID-19 are the reporting of successful therapeutical use of monoclonal antibodies anti-IL-6 receptors in COVID-19, whose use has been proved to be effective in several autoimmune diseases [3,4].

In this context polymorphisms at *IL6, IL6R* and *IL6ST* genes become interesting candidates to be prognostic and predictive markers on COVID-19 severity. IL-6 pathways have been reported to modulate the expression of *IL6* and *IL6R* genes, respectively [5] and to be associated to several inflammatory, autoimmune and infectious diseases [6–8]. Moreover, *IL6ST* gene mutations can cause Hyper-IgE recurrent infection syndrome-4B, a rare recessive immunologic disorder [9], highlighting *IL6ST* as a promising candidate gene on COVID-19.

Hence, the present study evaluated the role of polymorphisms at *IL6, IL6R* and *IL6ST* in COVID-19 severity aspects, along with their influence in IL-6 plasma levels during active COVID-19.

## Material and Methods

This is a cross-sectional study approved by the National Research Ethics Committee Ministry of Health e Ethics Committee in Research with Human Beings of the Institute of Biology Federal University of Pará (CAAE: 31800720.1.0000.0018). The sample is constituted by 227 patients collected during the period from September 2020 to July 2021, being considered as positive for COVID-19 those with positive RT-PCR or antigen test. Only patients unvaccinated at any time point of the study were considered for sampling.

Demographic and clinical data were collected from patients, including gender, age, and COVID-19-specific data (date of onset of symptoms, symptoms at diagnosis of COVID-19, hospitalization, need for oxygen and comorbidities). Patients were grouped according hospitalization.

Genomic DNA was isolated from blood samples using commercial kit (Wizard® Genomic DNA Purification Kit; Promega), following the fabricant protocol. Genotyping of rs1800795 (4351376 C 1839697_20), rs2228145 (4351376 C 16170664_10) and rs7730934 (4351379 C 3248953_10), localized at genes *IL6, IL6R* and *IL6ST*, respectively, was performed by customized TaqMan® assays, commercially provided by ThermoFisher. The real-time PCR was carried out in StepOnePlus™ Real-Time PCR System, following standard protocols.

Cytokine plasma levels have been previously evaluated in this sample [10] and IL-6 levels in the samples of the present study were retrieved and used for analysis. From 227 samples used in this study 95 were collected during active infection in hospitalized patients and used to correlate genotypes with IL-6 levels.

### Statistical strategies and tests

Two groups of patients were considered, hospitalized and non-hospitalized. The allele and genotype frequencies were obtained by direct count and the genotype frequencies were tested for Hardy-Weinberg equilibrium. Genotypic frequencies between groups were tested by Fisher exact test.

A control group, constituted by *IL6* genotype and allele frequencies representative of the same population (city of Belém), was used to comparison with COVID-19 subsamples. The frequencies of 300 individual were retrieved from a study performed before the pandemics [11]. IL-6 levels between different genotypes were compared by Mann-Whitney test.

The frequency of symptoms between hospitalized and non-hospitalized was compared by Wilcoxon paired test. Additionally, the frequency of symptoms was also compared between genotypes of all three SNPs.

## Results

Clinical and demographic characteristics of the sample is presented in Table 1, while the Table 2 presents the genotype and allele frequencies in the sample and subsamples.

**Table 1.** Demographic, epidemiological and clinical characterization of the sample.

| Variables | Not hospitalized<br>n= 95 (41,85%) | Hospitalized<br>n= 132 (58,15%) |
| --- | --- | --- |
| **Sex** |  |  |
| Women | 49 (51,58) | 58 (43,94) |
| **Age (years)** |  |  |
| 18-39 | 50 (52,63) | 29 (21,97) |
| 40-59 | 42 (44,21) | 67 (50,76) |
| ≥60 | 3 (3,16) | 36 (27,27) |
| Average | 39 | 51 |
| **Comorbidities** |  |  |
| Yes | 15 (14,79) | 54 (40,91) |
| **Ventilatory Support** |  |  |
| No | 95 (100) | 60 (45,45) |
| Non-invasive | 0 (0,0) | 70 (53,03) |
| Invasive | 0 (0,0) | 2 (1,51) |
| **Symptoms** |  |  |
| Fever | 58 (61,05) | 99 (75) |
| Cough | 46 (48,42) | 108 (81,81) |
| Coryza | 38 (40) | 47 (35,61) |
| Headache | 58 (61,05) | 71 (53,79) |
| Sore throat | 40 (42,10) | 43 (32,57) |
| Chest pain | 30 (31,58) | 56 (42,42) |
| Abdominal pain | 16 (16,84) | 31 (23,48) |
| Myalgia | 51 (53,68) | 79 (59,85) |
| Nausea and/or vomiting | 17 (17,89) | 45 (34,09) |
| Diarrhea | 33 (34,74) | 63 (47,73) |
| Dyspnea | 28 (29,47) | 80 (60,61) |
| Weakness or fatigue | 19 (20) | 36 (27,27) |
| Anosmia and/or ageusia | 60 (63,16) | 52 (39,39) |

n=number of patients.

**Table 2.**
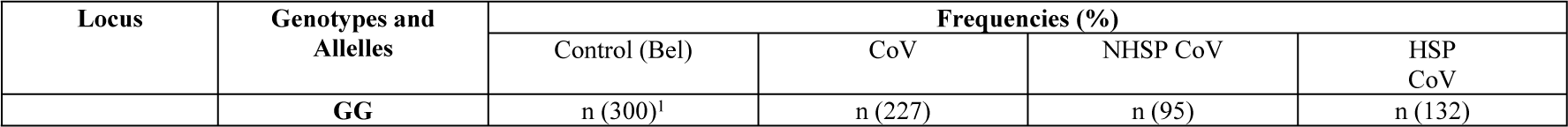
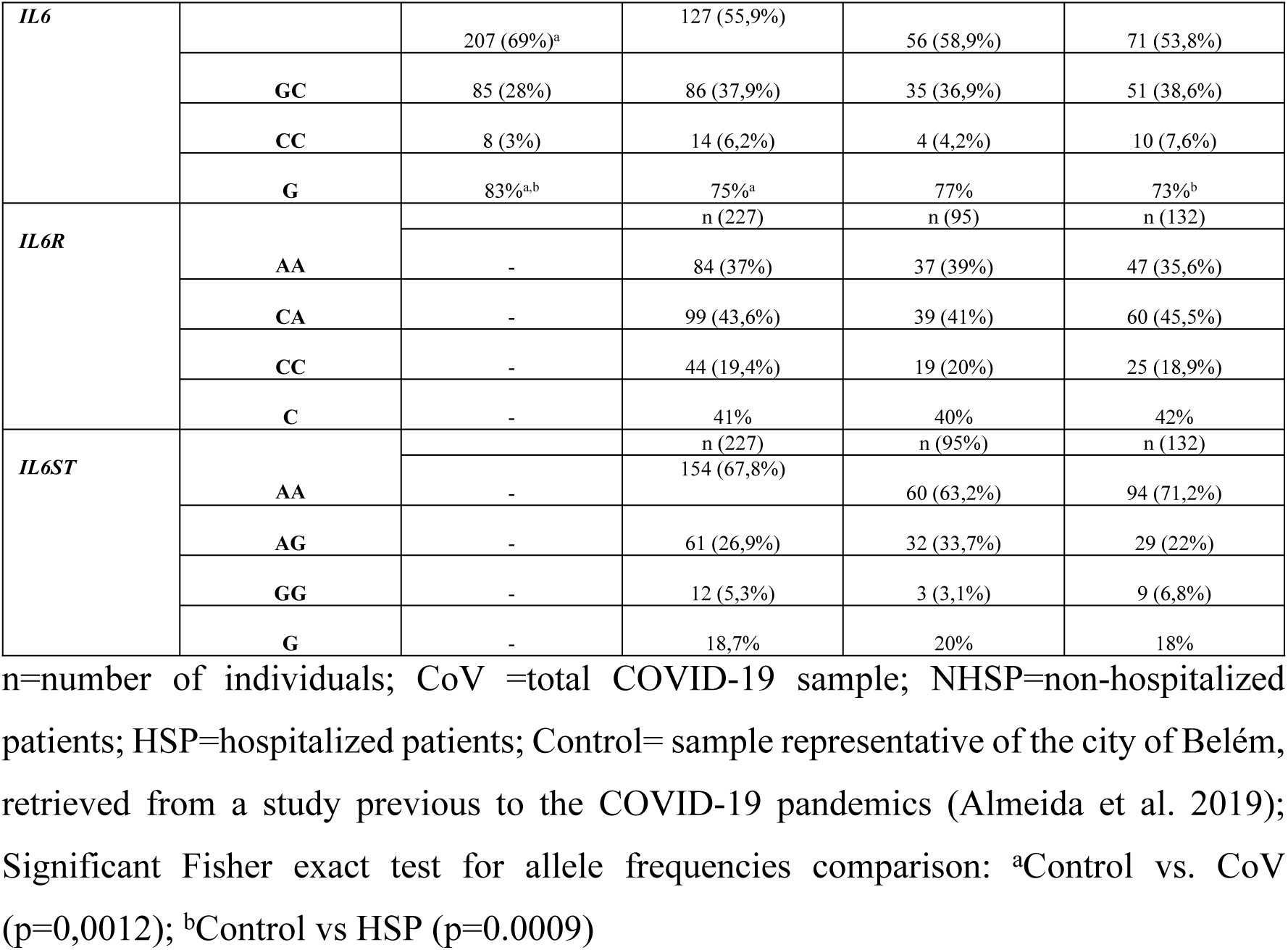
Genotype and allele frequencies of rs1800795, rs2228145 and rs7730934, localized at genes *IL6, IL6R* and *IL6ST*, respectively.

The Table 3 presents the frequencies of COVID-19 symptoms according SNP genotypes.

**Table 3.** Frequency of major COVID-19 symptoms according *IL6, IL6R* and *IL6ST* genotypes.

| **Locus** | **IL6R** |  | **IL6** |  | **IL6ST** |  |
| --- | --- | --- | --- | --- | --- | --- |
| **Genotype** | **AA** | **CC** | **GG** | **CC** | **AA+AG** | **GG** |
| **N** | 84 | 44 | 127 | 14 | 72 | 151 |
| **Fever** | 73.81 | 65.91 | 66.14 | 85.71 | 68.06 | 68.87 |
| **Cough** | 69.05 | 77.27 | 63.78 | 50 | 62.5 | 69.54 |
| **Coryza** | 32.14 | 40.91 | 36.22 | 35.71 | 36.11 | 37.09 |
| **Retroocular pain** | 25 | 31.82 | 21.26 | 50 | 25 | 28.8 |
| **Headache** | 59.52 | 61.36 | 54.33 | 64.29 | 48.61 | 60.26 |
| **Sore throat** | 39.29 | 36.36 | 28.35 | 42.86 | 27.78 | 40.4 |
| **Chest pain** | 36.9 | 47.73 | 33.07 | 42.86 | 40.28 | 37.75 |
| **Abdominal pain** | 27.38 | 22.73 | 17.32 | 28.57 | 25 | 19.21 |
| **Myalgia** | 60.71 | 63.64 | 51.18 | 57.14 | 54.17 | 58.28 |
| **Nausea** | 22.62 | 36.36 | 22.05 | 50 | 19.44 | 26.49 |
| **Vomiting** | 13.1 | 11.36 | 11.81 | 7.14 | 11.11 | 11.26 |
| **Diarrhea** | 45.24 | 45.45 | 38.58 | 42.86 | 44.44 | 41.06 |
| **Dyspnea** | 46.43 | 56.82 | 42.52 | 64.29 | 50 | 46.36 |
| **Weakness** | 51.19 | 61.36 | 46.46 | 42.86 | 48.61 | 54.97 |
| **Fatigue** | 57.14 | 65.91 | 55.12 | 64.29 | 51.39 | 62.91 |
| **Anosmia** | 45.24 | 50 | 46.46 | 57.14 | 48.61 | 47.68 |
| **Ageusia** | 44.05 | 54.55 | 43.31 | 64.29 | 48.61 | 45.7 |

N= sample size.

Comparison performed between genotypes revealed that carriers of the genotype *IL6R* CC have higher frequency of symptoms than AA carriers (Wilcoxon paired test; Z=2,6; p=0,0092); *IL6* CC carriers have higher frequency of symptoms than AA carriers (Z=2,72; p=0,0056). no differences were observed between *IL6ST* genotypes. (p=0.084).

The retrieving of data on IL-6 plasmatic levels from our previous published study [10] showed that among 95 patients with active COVID-19 the levels of plasmatic IL-6 was higher among *IL6* CC genotype carriers (seven patients; average 40,11 pg/ml) than among GG carriers (54 patients; average 31,2 pg/ml), being this difference statistically significant (Mann-Whitney test; Z(U)=-1,98; p=0,04).

## Discussion

In general, our data agrees with trends from literature in relation to higher proportions of males, elder patients and comorbidity carriers in the hospitalized subsample [12–13]. In relation to symptoms taste/smell was more frequently observed in non-hospitalized patients, corroborating literature reports that associate these symptoms with milder COVID-19 [14].

The association of the allele C of the rs1800795, localized at *IL6* gene, agrees with previous meta-analysis published by our group that correlated the frequency of the C allele with the COVID-19 mortality rates worldwide [15].

The association of the *IL6* genotype CC is also corroborated by the higher frequency of COVID-19 symptoms among CC carriers (Table 3), as well as by the slight increase of IL-6 levels in CC in comparison to GG genotypes, during the active disease.

The understanding of the functional basis of association between *IL6* C allele with COVID-19 severity is still not clear. The role of this variant in *IL6* expression is not conclusive, there are a trend toward higher IL-6 production among the allele C carriers as well as an association of this allele with more sever forms of pneumonia in general, as previously suggested in a recent meta-analysis [16]. In this context an additional association of the *IL6R* genotype CC with higher frequencies of symptoms could be also observed in our results.

While the association of *IL6* genotypes with serum IL-6 levels is still not completely understood, the *IL6R* genotype CC is more consistently associated with higher plasmatic levels of this receptor [5,17]. In particular, the plasmatic IL-6R is related to the trans-activating pathway, likely associated with proinflammatory response and chronic inflammatory diseases [18]. This proinflammatory trend is also corroborated by the successful use of IL-6R blockade by monoclonal antibodies in some inflammatory diseases [19]. Hence, some studies claim that the blockade of IL-6R could be beneficial in COVID-19 therapy protocols and even considering the influence of IL6R [3] polymorphism in these protocols [20].

Thus, the higher frequency of COVID-19 symptoms among *IL6R* CC genotypes could reflect the higher expression profile of this genotype and its consequent proinflammatory effects.

In conclusion our results point to a consistent association of *IL6* and *IL6R* polymorphisms with COVID-19 severity, putatively due to higher expression of these genes related to CC genotypes and their proinflammatory consequences, corroborating previous meta-analysis studies correlating them with mortality rates worldwide and their role in pneumonia and immunobiological therapeutic protocols involving IL-6 pathways.

## Funding

The study was supported by the National Council for Scientific and Technological Development (CNPQ #401235/2020-3); Amazon Foundation for Research Support (FAPESPA#060/2020).

## Acknowledgments

The authors thank all patients who agreed to voluntarily participate in this study.

## Notes

### Competing Interest Statement

The authors have declared no competing interest.

## References

1. Ladds E, Rushforth A, Wieringa S, Taylor S, Rayner C, Husain L, et al. Persistent Symptoms after Covid-19: Qualitative Study of 114 ‘Long Covid’ Patients and Draft Quality Criteria for Services. BMC Health Services Research. 2020; 1–13. doi: 10.1101/2020.10.13.20211854.

2. Wolf J, Rose-John S, Garbers C. Interleukin-6 and Its Receptors: A Highly Regulated and Dynamic System. Cytokine. 2014; 70 (1): 11–20. doi: 10.1016/j.cyto.2014.05.024.

3. Veerdonk F L, Giamarellos-Bourboulis E, Pickkers P, Derde L, Leavis H, van Crevel R, et al. A Guide to Immunotherapy for COVID-19. Nature Medicine. 2022; 28 (1): 39–50. doi: 10.1038/s41591-021-01643-9.

4. Lim P C, Wong K L, Rajah R, Chong M F, Chow T S, Subramaniam S, et al. Comparing he Efficacy of Tocilizumab with Corticosteroid Therapy in Treating COVID-19 Patients: A Systematic Review and Meta-Analysis. DARU, Journal of Pharmaceutical Sciences. 2022; 30 (1): 211–28. doi: 10.1007/s40199-021-00430-8.

5. Smith J P A, Humphries S E. Cytokine and Cytokine Receptor Gene Polymorphisms and Their Functionality. Cytokine and Growth Factor Reviews. 2009; 20 (1): 43–59. doi: 10.1016/j.cytogfr.2008.11.006.

6. Jones S A, Hunter C A. IL-6 as a Keystone Cytokine in Health and Disease. Nature Immunology. 2015; 16 (5): 448–57. doi: 10.1038/ni.3153.The.

7. Mihara M, Hashizume M, Yoshida H, Suzuki M, Shiina M. IL-6/IL-6 Receptor System and Its Role in Physiological and Pathological Conditions. Clinical Science. 2012; 122 (4): 143–59. doi: 10.1042/CS20110340.

8. Rose-John S, Winthrop K, Calabrese L. The Role of IL-6 in Host Defence against Infections: Immunobiology and Clinical Implications. Nature Reviews Rheumatology. 2017; 13 (7): 399–409. doi: 10.1038/nrrheum.2017.83.

9. Schwerd T, Twigg S R F, Aschenbrenner D, Manrique S, Miller K A, Taylor I B, et al. A Biallelic Mutation in IL6ST Encoding the GP130 Coreceptor Causes Immunodeficiency and Craniosynostosis. Journal of Experimental Medicine. 2017; 214 (9): 2547–62. doi: 10.1084/jem.20161810.

10. Queiroz M A F, Neves P F M, Lima S S, Lopes J C, Torres M K S, Vallinoto I M V C, et al. Cytokine Profiles Associated With Acute COVID-19 and Long COVID-19 Syndrome. Frontiers in Cellular and Infection Microbiology. 2022; 12 (June): 922422. doi: 10.3389/fcimb.2022.922422.

11. Almeida N C C, Queiroz M A F, Lima S S, Costa I B, Fossa M A A, Vallinoto A C R, et al. Association of Chlamydia Trachomatis, C. Pneumoniae, and IL-6 and IL-8 Gene Alterations with Heart Diseases. Frontiers in Immunology. 2019; 10 (FEB): 1–14. doi: 10.3389/fimmu.2019.00087.

12. Wu G, Yang P, Xie Y, Woodruff H C, Rao X, Guiot J, et al. Development of a Clinical Decision Support System for Severity Risk Prediction and Triage of COVID-19 Patients at Hospital Admission: An International Multicentre Study. European Respiratory Journal. 2020; 56 (2). doi: 10.1183/13993003.01104-2020.

13. Li X, Xu S, Yu M, Wang K, Tao Y, Zhou Y, et al. Risk Factors for Severity and Mortality in Adult COVID-19 Inpatients in Wuhan. Journal of Allergy and Clinical Immunology. 2020; 146 (1): 110–18. doi: 10.1016/j.jaci.2020.04.006.

14. Gonçalves L F, Gonzáles A I, Paiva K M, Patatt F S A, Stolz J V, Haas P. Smell and Taste Alterations in COVID-19 Patients: A Systematic Review. Revista Da Associacao Medica Brasileira. 2021; 66 (11): 1602–8. doi: 10.1590/1806-9282.66.11.1602.

15. Leite M M, Gonzalez-Galarza F F, Silva B C C, Middleton D, and Eduardo José Melo dos Santos. Predictive Immunogenetic Markers in COVID-19. Human Immunology 2021; 82 (4): 247–54. doi: 10.1016/j.humimm.2021.01.008.

16. Ulhaq Z S, Soraya G V. Anti-IL-6 Receptor Antibody Treatment for Severe COVID-19 and the Potential Implication of IL-6 Gene Polymorphisms in Novel Coronavirus Pneumonia. Medicina Clinica. 2020; 155 (12): 548–56. doi: 10.1016/j.medcli.2020.07.002.

17. Galicia J C, Tai H, Komatsu Y, Shimada Y, Akazawa K, Yoshie H. Polymorphisms n the IL-6 Receptor (IL-6R) Gene: Strong Evidence That Serum Levels of Soluble IL-6R Are Genetically Influenced. Genes and Immunity. 2004; 5 (6): 513–16. doi: 10.1038/sj.gene.6364120.

18. Scheller J, Garbers C, Rose-John S. Interleukin-6: From Basic Biology to Selective Blockade of pro-Inflammatory Activities. Seminars in Immunology. 2014; 26 (1): 2–12. doi: 10.1016/j.smim.2013.11.002.

19. Jones S A, Jenkins B J. Recent Insights into Targeting the IL-6 Cytokine Family in Inflammatory Diseases and Cancer. Nature Reviews Immunology. 2018; 18 (12): 773–89. doi: 10.1038/s41577-018-0066-7.

20. Garbers C, Rose-John S. Genetic IL-6R Variants and Therapeutic Inhibition of IL-6 Receptor Signalling in COVID-19. The Lancet Rheumatology. 2021; 3 (2): e96–97. doi: 10.1016/S2665-9913(20)30416-1.

